# Enrichment of phenotype among biological forms of *Anopheles stephensi* Liston through establishment of isofemale lines

**DOI:** 10.1101/2022.09.28.509862

**Authors:** Chaitali Ghosh, Naveen Kumar, Raja Babu Singh Kushwah, M Soumya, Soumya Gopal Joshi, R Chethan Kumar, Tejashwini Alalamath, Subhashini Srinivasan, Suresh Subramani, Sampath Kumar, Sunita Swain

## Abstract

The success of vector management programs relies on knowledge of the biology and genetic make-up of mosquitoes so that they can be interlaced with modern tools for developing suitable intervention strategies. There are many reports available for rearing varied species of mosquito vectors. However, there are limited studies addressing the development of isofemale lines among mosquitoes to homogenize the population to obtain both high-quality genome assemblies and enrichment of phenotype. *Anopheles stephensi*, an urban malaria vector, is one of the major invasive vectors of malaria distributed throughout the Indian subcontinent, Middle East, and has recently been expanding its range in Africa. With the existence of three biological forms, distinctly identifiable based on the number of ridges on eggs with varying vectorial competence, *An. stephensi* is a perfect species for developing a method for the successful establishment of isofemale lines, which can be tested for retention of the expected vectorial competence for the various forms. We describe the key steps in the establishment and validation of isofemale lines, which include monitoring the transgenerational fitness traits, morphometrics of eggs, and adult wing size during every generation. After the initial inbreeding depression, as proof of the tedious selection process, no significant morphometric differences were observed in the wings and egg size between the parental and their respective isofemale lines. We observed a significant change in the vectorial competence between the respective isofemale and parental lines enriching expected differential susceptibility towards malaria parasites by the type and intermediate forms. Interestingly, IndCh and IndInt strains showed variations in resistance to different insecticides belonging to all the four major classes. These variant lines have been characterized for their levels of homozygosity both at the phenotype and genotype levels and can be used as a standard reference or as a biological resource for other studies related to urban malaria research.

**Author summary:** Isofemale lines can be a valuable resource for characterizing and enhancing several genotypic and phenotypic traits. This is the first detailed report of the establishment of two isofemale lines of type and intermediate biological forms in *Anopheles stephensi*. The work encompasses the characterization of fitness traits among the two lines through a transgenerational study. Further, isofemale colonies were established and used to characterize insecticide susceptibility and vector competence. The study provides valuable insights into the differential susceptibility status of the parental and isofemale to different insecticides belonging to the same class. Corroborating with the earlier hypothesis we exemplify the high vector competence in type form than the intermediate from using homozygous isofemale lines. Using these isofemale lines it is now possible to study host-parasite interactions and identify factors that might be responsible for altered susceptibility and increased vector competence in *An. stephensi* biological forms that would also pave way for developing better vector management strategies.

## Introduction

The adaptive evolution of mosquitoes dates back to the dinosaurs from the Mesozoic era to species in the Holocene era (considered as the deadliest animal) [1]. Owing to the diversity of mosquitoes and their rapid rate of evolution [2–4], many research groups across the world work on various aspects of the mosquito vector ranging from its biology, molecular genetics, physiology, population genetics, developmental biology, evolutionary biology, behaviour, and so on. During these studies, colonization of mosquitoes under laboratory conditions for the purpose of experimentation becomes essential. Over recent years several researchers have developed techniques to rear mosquito species that were previously considered difficult to maintain under laboratory conditions [5, 6]. One such effort led to the colonization of *Anopheles stephensi* under controlled environmental conditions. The rearing protocols with optimal temperature, light, and humidity are established for standard insectary operations [7–9]. *An. stephensi* exhibits a greater magnitude of variation across populations, demanding the establishment of isofemale lines with greater genetic homogeneity, enabling the study of the quantitative and qualitative traits under laboratory conditions [10]. The molecular variations at genomic, transcriptomic, and proteomic levels can be better established in isofemale lines [11]. Isofemale lines have been previously used for studying behavior [12], morphometry [13], insecticide resistance [14, 15], pathogen susceptibility, and speciation among mosquitoes and other insect species. By enrichment for phenotypes, isofemale lines enable the identification of associated genotypes, which are otherwise diluted in their heterozygous parent lines [16]. Establishing an isofemale line has been shown to delineate the non-additive genetic variance due to local adaptations, bottleneck events, and other epistatic effects [17].

Furthermore, when isofemale lines are maintained as isofemale colonies for many generations, they can be used for any live material experimental studies [18]. During the initial stages of colonization, insect populations pass through a genetic bottleneck [19]. Quantitative traits tend to vary greatly due to adaptation to the rearing conditions. However, since insectary operations are well standardized, maintaining the isofemale lines under these regimes would not deviate from the quantitative traits determined from the mean values. Though there are publications related to the establishment of isofemale lines, the technique and success vary depending on the species (*Drosophila* [17], *Trichogramma* [20], *Chilo* sp. [18], and mosquitoes [13, 21]) in the insect kingdom depending on their biology. There are many instances of unsuccessful establishment of isofemale lines due to improper mating, poor reproductive success, inbreeding depression, etc. [22]. Among many hurdles, inbreeding depression is a daunting challenge that can be mitigated with careful line establishment practices and selection of the founder females in the filial generation with key fitness traits [13, 20, 23, 24]. In the present paper, we outline a detailed procedure for the establishment of two isofemale lines from insectary colonized populations. A type form “IndCh” derived from TIGS-2 (T2) and an intermediate form “IndInt”, derived from TIGS-6 (T6) [25]. We report the characterization of the following parameters: (a) fitness, (b) homozygosity across filial generations, (c) insecticide susceptibility, and (d) vectorial competence for both these lines.

## Materials and Methods

### Collection and colonization of mosquitoes

Mosquitoes were collected from their natural habitats through larval or adult sampling as per the previously established protocols [26]. The T6 strain was originally collected from the semi-urban area of Sriramanahalli, Bengaluru, Karnataka (12.972442°N, 77.580643°E) at the larval stage in 2016. The T2 strain was collected from the urban area of Anna Nagar, Chennai, Tamil Nadu (13.018410°N, 80.223068°E) at the adult stage in 2016. The rearing protocols were standardized as per previously established protocols [27]. The adults were identified using the identification keys of Nagpal and Sharma [28]. On day 6 or 7, the adults were blood-fed through a standard membrane feeding system (approval Ref. No. TIGS 2nd IBSC Oct 2018).

### Establishment of isofemale lines

The isofemale progenies were developed by improvising a method previously described by Ghosh and Shetty [29]. About 20-25 gravid females (G0) were separated from their parental colonies (T2 and T6) maintained in the TIGS insectary. On day 3, each gravid female was transferred carefully to a single ovicup for egg laying. The individual females were allowed to lay eggs undisturbed for ~24 to 48 h under laboratory conditions. The single G0 female that laid the highest number of eggs and had highest percent hatchability was selected for the establishment of isofemale lines. Eggs collected from the single female were checked for egg ridge number and allowed to hatch. Larvae were provided with larval food and reared to adults as described earlier. Emerged adult siblings (G1) were kept in a rearing cage and allowed to inbreed (sibling mating). Around 5-10 mated females were randomly separated for continuing the filial generations. The same protocol was followed in subsequent filial generations (20 generations for IndCh and 23 generations for IndInt). Key life-table parameters like fecundity (number of eggs produced per female), egg morphology, percent egg hatchability, number of larvae and pupae, sex ratio (male: female), longevity of adult females (lifespan), and frequency of blood meals taken by a female, were recorded in every generation to check the fitness of the isofemale lines.

### Establishment of isofemale colony

Both IndCh and IndInt isofemale lines were maintained as two separate isofemale colonies after 20 generations and 23 generations respectively, instead of single female progeny [29]. These two lines are being maintained at present in TIGS insectary. In both the colonies, males and females were allowed to inbreed within the filial generation, blood-fed, and allowed to lay eggs in masses. All the key fitness parameters were monitored and recorded. These two isofemale colonies were utilized for insecticide-susceptibility assays and vectorial competence studies with *Plasmodium berghei* (*P. berghei*) and *Plasmodium falciparum* (*P. falciparum*).

### Morphometric analyses

Different developmental stages of the mosquitoes were carefully measured for their morphometric characteristics. In the present study, we have measured the eggs and wings in parental and isofemale lines.

#### • Egg parameters

In total, 5 parameters viz. egg ridge number, shape, length and width of egg, and egg float from randomly selected 15 individuals were analysed. Eggs were placed on a wet filter paper and measured under the microscope with an ocular micrometer (Unilab GE-34, Binocular Research Microscope, India) and recorded. The egg shape, ridge numbers, and floaters were studied following earlier protocols [30, 31]. In isofemale lines, egg ridge was counted in every alternate generation.

#### • Measurement of wing length

A total of 15 wing samples of both male and female mosquitoes were selected for wing measurement. The wings of male and female mosquitoes were carefully dissected 10 days after eclosion. The mosquitoes were anesthetized on CO_2_ pads and the right wing dissected under a microscope (Olympus, SZX2-ILLK, Germany). Additional care was taken to ensure the wings were not damaged or folded during mounting. The wings were measured from the distal end of the alula to the tip between R1 and R2 veins, excluding the fringe scale as they are normally curved or wither off during mounting [32]. Wings were photographed and recorded under a stereomicroscope (Leica MZ10F, Germany). The photographed wings were measured using Leica Application Suite X (LAS X) software. The measurements were carried out separately by two observers to minimize error.

### Estimation of homozygosity

The Fastq files for the 3 reference genomes were downloaded from NCBI SRA (Sequence read archive). The 3 genomes (IndCh = SRR15146350, IndInt = SRR15603373, STE2 = SRR1168951) were mapped against the UCI 2.0 reference genome using bowtie2 to call variants and generate a VCF file. The VCF was filtered for DP=3, QUAL=10, and Chr 2, 3, X, and MAF (Minimum allele frequency) = 0.01 and merged. The merged VCF file was converted to a matrix file containing numeric genotype information (0,1,2) representing the three possible genotype for each SNP. The information was used in the table to quantify heterozygosity–homozygosity in the genomes.

### Insecticide-susceptibility assays

Five-day-old adult female mosquitoes of the parental and isofemale colonies were used for insecticide-susceptibility assays as per WHO guidelines. Insecticide-impregnated papers of discriminating doses and insecticide resistance monitoring test kit ordered from Vector Control Research, Universiti Sains Malaysia (a WHO referral Centre for these kits and impregnated papers) [33].

Each replicate consisted of 25 females and four such replicates were used for the experiment. One corresponding control was tested with the control papers supplied with the kit for respective classes of insecticides. They were exposed to eight insecticides representing all four major insecticide classes (Table 1). Mosquitoes used for DDT, permethrin, and deltamethrin were pre-exposed to PBO (synergist) to understand the role of the metabolic mechanism of resistance. Assays were performed with 1h exposure followed by 24h recovery time, where the mosquitoes were provided with 10% glucose solution soaked on cotton pads. The mortality was recorded 24h post-exposure and all the mosquitoes (dead and alive tested) were kept in individual microfuge tubes and preserved at −20°C for future analysis. All the susceptibility assays were carried out at the TIGS insectary at 27±1°C and 75±5% RH.

**Table 1:** List of insecticides, their respective class and concentrations used for the insecticide-susceptibility assays.

| Sl. No. | Insecticide | Class | Discriminating concentration | Exposure time |
| --- | --- | --- | --- | --- |
| 1 | DDT | Organochlorines | 4% | 1 h |
| 2 | Dieldrin |  | 4% |  |
| 3 | Malathion | Organophosphates | 5% |  |
| 4 | Fenitrothion |  | 1% |  |
| 5 | Bendiocarb | Carbamates | 0.1% |  |
| 6 | Propoxur |  | 0.1% |  |
| 7 | Permethrin | Synthetic pyrethroids | 0.75% |  |
| 8 | Deltamethrin |  | 0.05% |  |
| 9 | PBO | (Synergist) | 4% |  |

### Vectorial competence studies

The vector competence studies were carried out in parental and isofemale colonies. The competence was evaluated in both in vivo and in vitro assays using the *P. berghei* and *P. falciparum* parasites.

#### • Plasmodium berghei

*Albino* mice (BALB/c) were provided by the Animal Care Resource Center (ACRC) of the National Centre for Biological Sciences (NCBS), Bengaluru, India. All animal procedures were operated and approved according to the Instem IAEC under project INS-IAE-2019/01(N). The rodent malarial parasite, *P. berghei* (ANKA strain-MRA-311, BEI resources, USA) was used for infecting mice by injection of 100–150 μl intraperitoneally per mouse [34]. Infected mice upon reaching 5–6% parasitemia and 1:1 or 1:2 male:female gametocyte ratio was anesthetized and exposed to 5-6 days old female mosquitoes as detailed below. Prior to keeping mice for feeding mosquitoes, the final count of parasitemia and exflagellation was recorded [34].

After eclosion, adult female mosquitoes were provided with 10% glucose solution for 4–6 days. Two replicate cups with 40 mosquitoes were kept fasting for 6 h prior to blood feeding. The mosquito cups were covered with a black cloth and left undisturbed for 30–40 min during blood feeding. The fully engorged females were separated from unfed and half-fed mosquitoes. The cups were maintained in a bioenvironmental chamber (Percival insect chamber I-30VL, Perry, USA) at 19°C and 75% RH (12:12h day and night cycles) for 13–14 days. A cotton wool pad soaked with 10% D-glucose; 0.05% para-aminobenzoic acid (PABA) solution was changed every alternate day until dissection. The mosquito midguts were dissected on the 14^th^ day using PBS and stained with 1% mercurochrome. Midguts were examined for the presence of oocysts and recorded under microscope (Nikon eclipse Ni-U upright microscope, Japan).

#### • Plasmodium falciparum

*P. falciparum* (PfNF54 Line E, procured from BEI resources (MRA-1000)) was revived for standard membrane feeding assay (SMFA). The culture was initiated at 0.15–0.2% asexual parasitemia and 5% hematocrit in a 10ml complete medium (RPMI-1640 with 6 g/l of HEPES, 50 mg/l of hypoxanthine, 2.5 g/l of sodium bicarbonate and 10% human serum) (ethical approval Ref. No. inStem/IEC-12/001, dated 20/03/2019). The cultures were maintained in an atmosphere of 5% O_2_, 5% CO_2_, and 90% N_2_ for 16–18 days with daily media change. For the mosquito feeding experiment, staggered cultures (14 and 17 days old) were selected based on their stage-V gametocytaemia and exflagellation activities (above 10 ex-flagellating centers per 40X field) and pooled [35].

During the day of SMFA, 4–6 day old female mosquitoes (40 nos., in duplicates) were placed in paper cups and maintained with 5% D-glucose overnight. Sugar cotton was removed 4 h prior to blood feeding and kept in dark till SMFA [36]. Mosquitoes were fed with mature stage-V *P. falciparum* NF54 gametocytes as mentioned earlier. The temperatures were maintained at 37°C during SMFA. After 30 min of blood-feeding, fully engorged mosquitoes were maintained in an environmental chamber set at 26°C and 75% RH (Percival insect chamber I-30VL, Perry, USA). Mosquitoes were provided daily with fresh wet cotton balls soaked in 10% D-glucose and 0.05% PABA solution. The midguts were dissected on the 9th day and stained with 1% mercurochrome, and the number of oocysts per midgut was recorded using a light microscope (Nikon eclipse Ni-U upright microscope, Japan).

### Statistical analyses

Descriptive and inferential statistical analysis has been carried out in the present study. Various life-table parameters like fecundity, hatchability, pupation percentage, eclosion percentage, and male: female ratio were recorded and presented in graphical forms. For eggs and wing measurements statistical t-test analysis for independent or correlated samples was performed using Vassar Stat software (http://vassarstats.net/). The mean and SEM were compared between populations. p-value >0.05 was considered non-significant for each parameter. Insecticide susceptibility was evaluated using the Prism GraphPad software package (version 9.0). The vectorial competence between the mosquito populations was analyzed using a non-parametric Mann–Whitney test using Prism GraphPad.

## Results

### Transgenerational fitness during sibling mating

In each generation, several key life-table parameters like fecundity, percent egg hatchability, number of larvae and pupae, sex ratio (male: female), longevity of adult females (lifespan), and frequency of blood meal were recorded. A greater number of females was selected to start the isofemale line as we expected mortality during the initial filial generations due to inbreeding [37, 38]. The average number of eggs was around 80–100 per individual female and fecundity of both the lines normalized around the 15^th^ generation. A similar trend was also observed in hatching percentage. Post normalization, the hatchability of both the isofemale lines improved to 85%. The development of larvae to pupae (pupation %) (Fig. 2C), followed by the transformation of the pupae to adults (eclosion %) (Fig. 2D), were affected during sibling mating. There were instances where sharp dips were observed owing to the mortality of the larvae and pupae. Most of the pupal mortality was observed during eclosion as also observed in other studies [39]. The male: female ratio was calculated in each generation. The findings suggest (Fig. 3A, B) that the ratios tend to get skewed during the first few filial generations and balances out after successive generations (1:1.5) [23]. It was observed that there was no significant change in the longevity of adult females (lifespan) or in their blood-feeding behavior.

**Figure 1:**
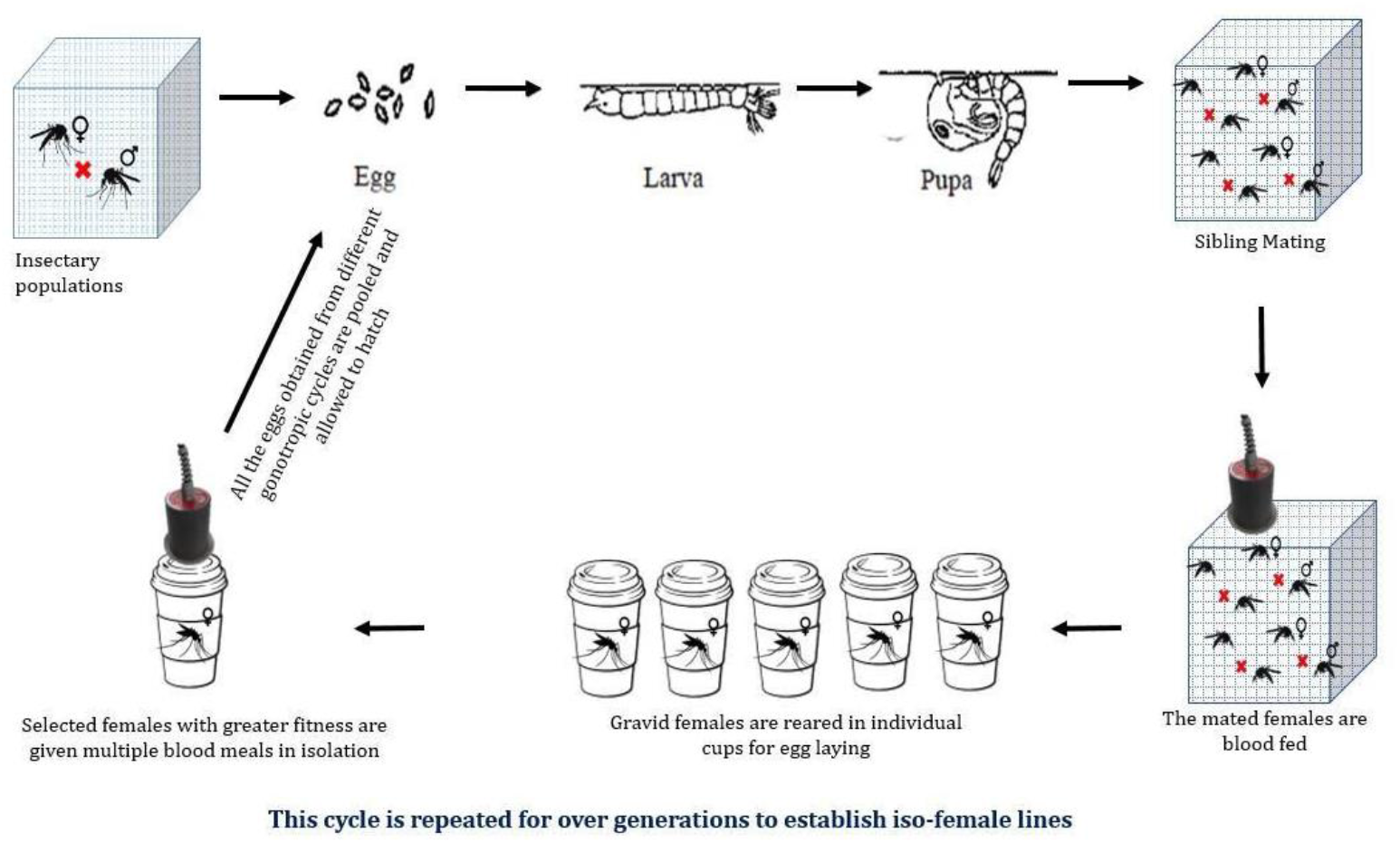
Schematic representation of the establishment of isofemale lines.

**Figure 2:**
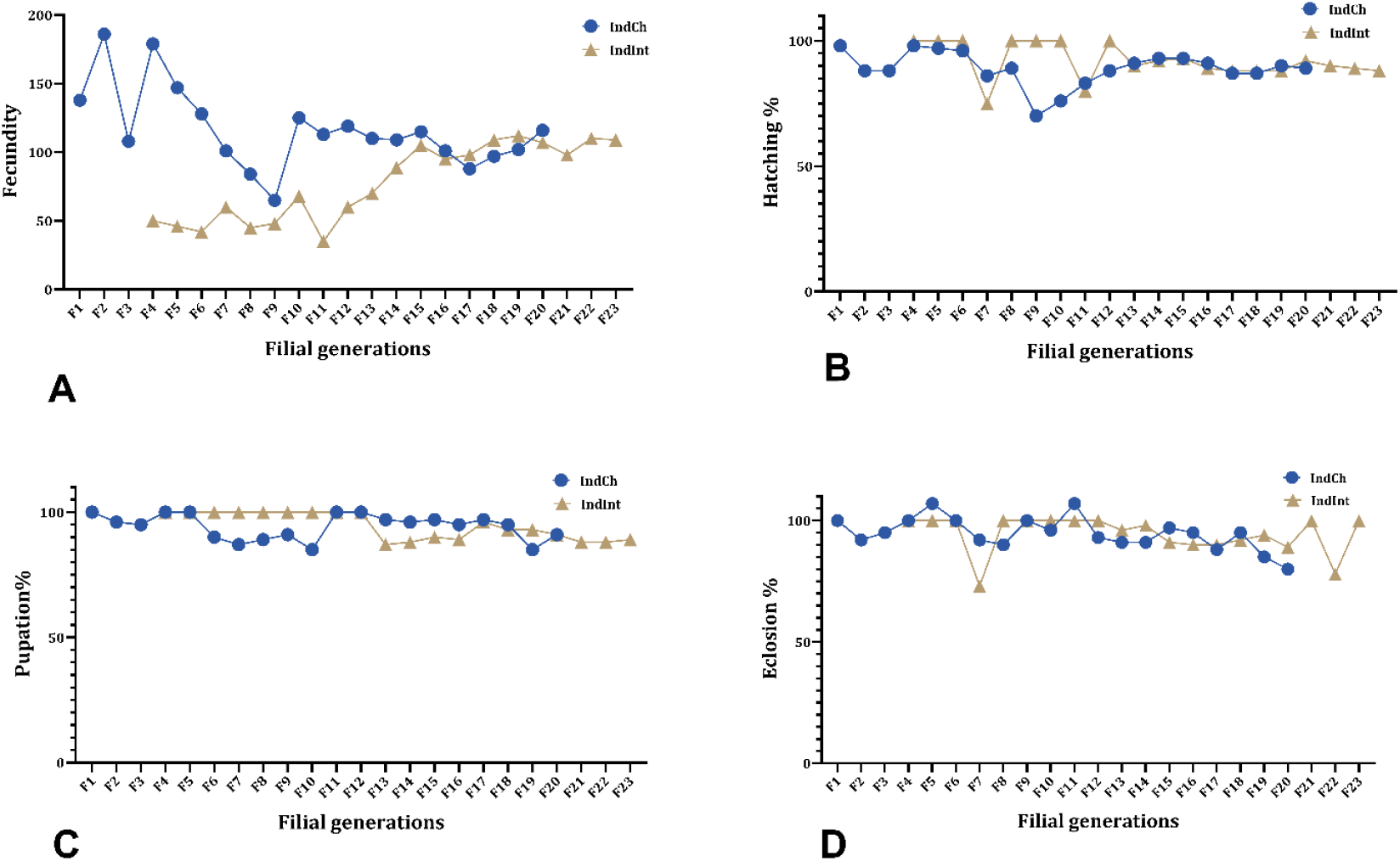
Graph of the two isofemale lines of *An. stephensi* (IndCh: F1-F20 generations and IndInt: F4–F23 generations) showing (A) transgenerational fecundity, (B) hatching %, (C) pupation %, and (D) eclosion % across filial generations.

**Figure 3:**
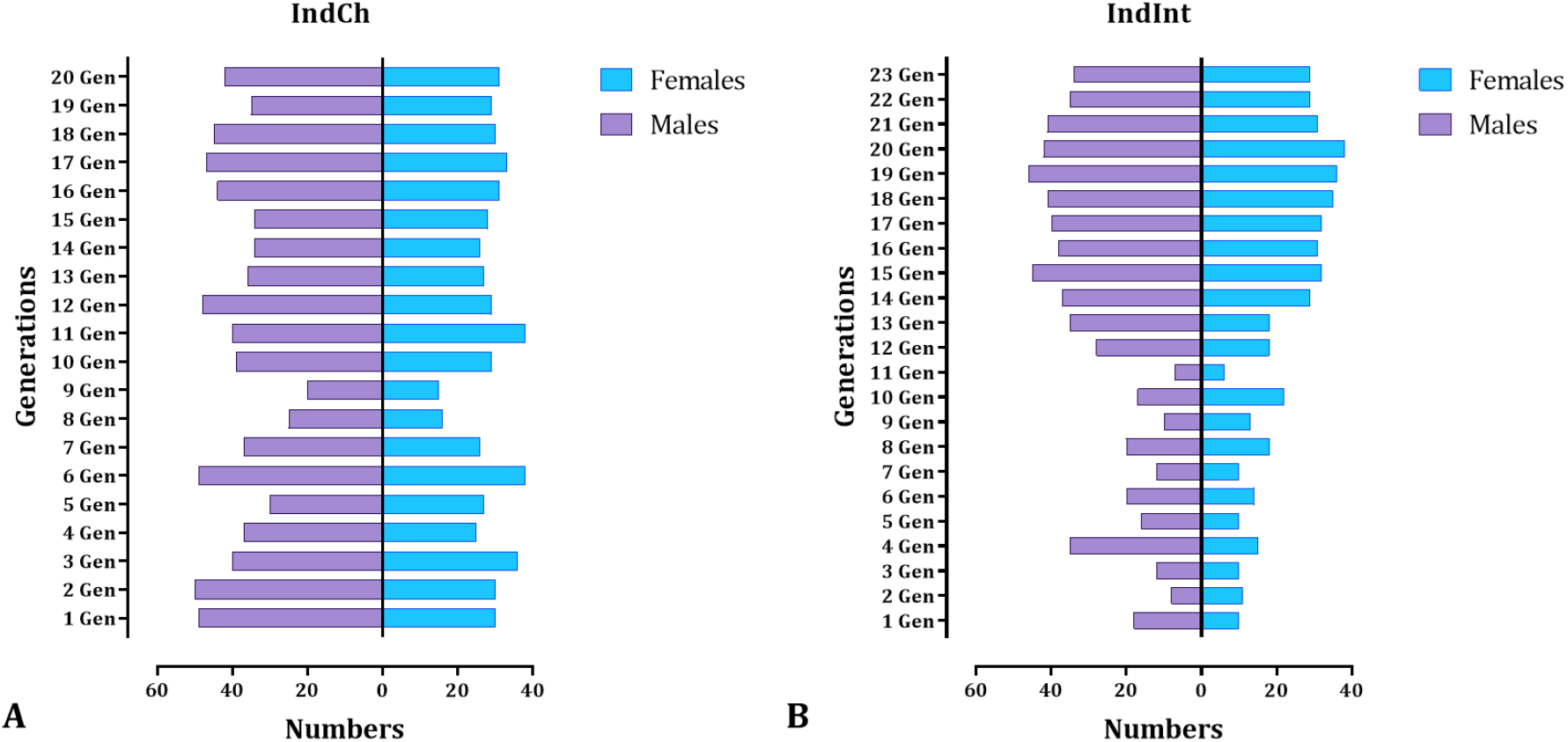
The male: female numbers among the two isofemale lines (A) IndCh isofemale and (B) IndInt isofemale line across filial generations (up to 20 and 23 generations, respectively).

### Morphometrics of egg and wings

To ascertain the morphometric changes that arise during and after sibling mating, several parameters such as egg shape, size, float numbers, float ridge, and wing length were observed and recorded (Table 2). The shape of the egg was slightly boat-shaped, black in color and blunt at anterior and posterior ends (Fig. 4) [60, 77]. The floats, which are an important morphological marker for differentiating the biological forms of different *Anopheles* species, showed minor variations in size. The maximum float lengths of IndCh and IndInt isofemale forms were 229.41±4.81 and 209.33±10.62 μm and the float widths were 71.17±6.96 and 69.33±7.98 μm, respectively. The float length-to-width ratios of IndCh and IndInt were 3.24±0.08 and 3.15±0.15, respectively. The maximum float lengths of IndCh and IndInt parental strains were 218.09±3.49 and 210.58±6.21 μm and the float widths were 71.17±6.96 and 69.33±7.98 μm, respectively. The float length-to-width ratios of IndCh and IndInt parental strains were 3.40±0.09 and 3.14±0.12, respectively (Table 2). The float size also influenced the number of ridges. For type and intermediate isofemale eggs, the number of float ridges were 20.81±0.08 (range 20–21) and 16.66±0.13 (range 16–17), respectively. For type and intermediate parental strains, it was 20.8±0.11 (range 19–21) and 17.35±0.31 (range 16–18). Many other parameters related to morphometric of egg are presented in Table 2. Overall, there were no significant differences in all the parameters of eggs when compared between the isofemale lines of IndCh (F=1.03, p>0.05) and IndInt (F=1.01, p>0.05), and also when compared with their respective parental lines.

**Figure 4A:**
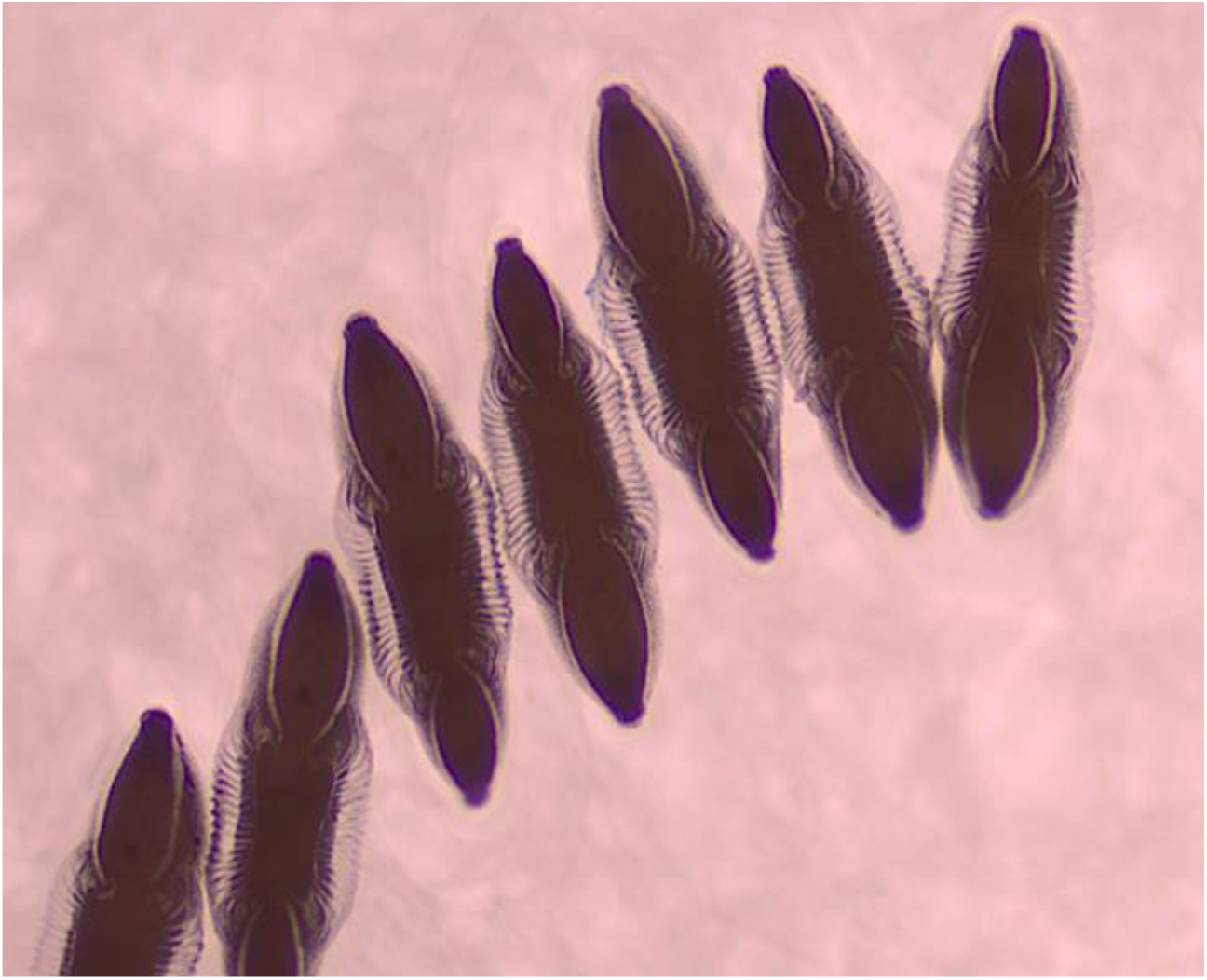
Eggs of IndCh of *An. stephensi* (magnification-10X)

**Figure 4B:**
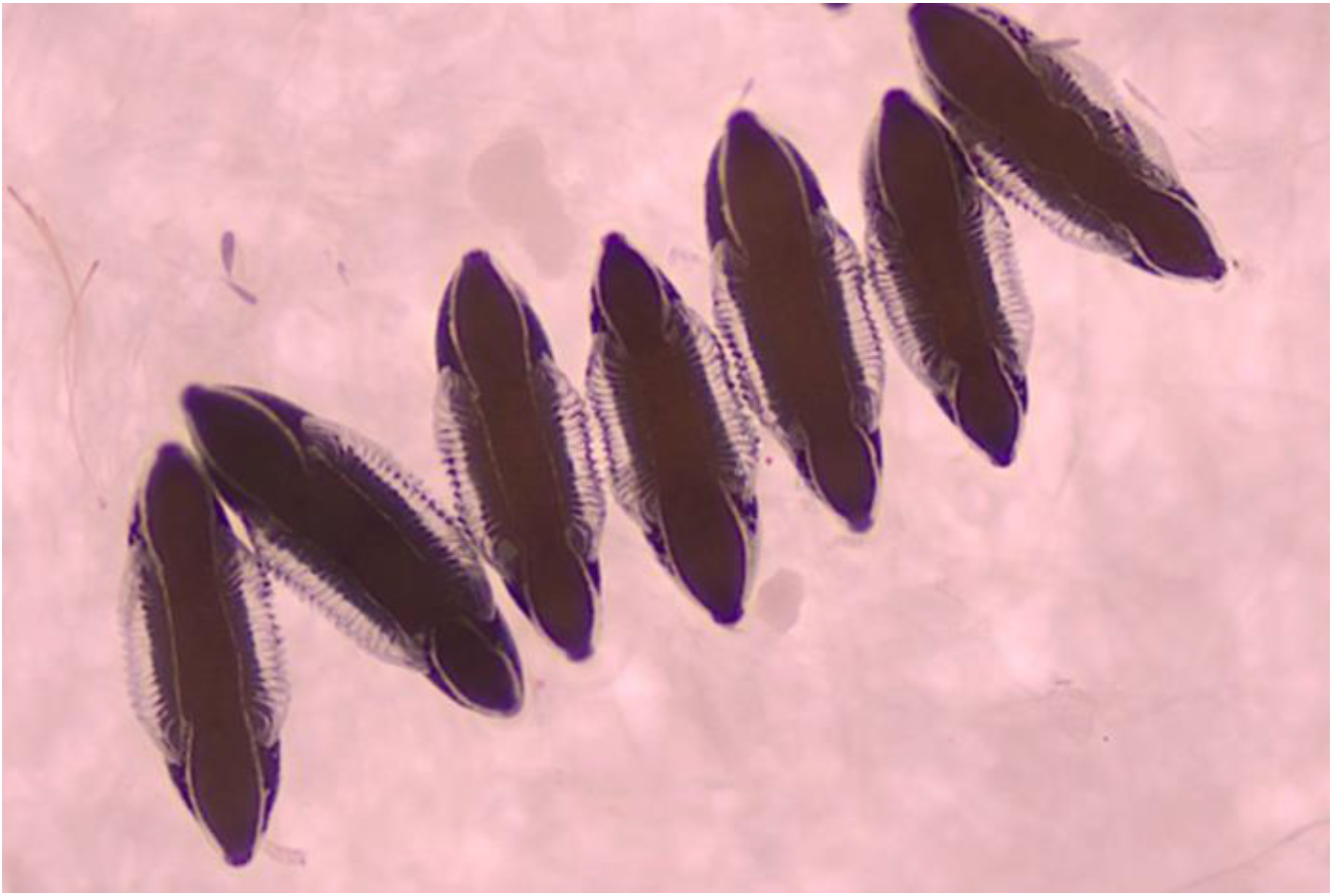
Eggs of IndInt of *An. stephensi* (magnification-10X)

**Table 2:** Egg measurement of paternal and isofemale strains of IndCh and IndInt biological forms. EL (length of egg), EW (width of egg), EL/EW (length/width of egg ratio), EFL (egg float length), EFW (egg float width), EFL/EFW (egg float length/egg float width ratio), FR (float ridge no.), EL/EFL (egg length ratio/egg float length), EFL/ER (egg float length/ egg ridge ratio), EFL/EW (egg float length/egg width ratio), EFW/ER (egg float width/ egg ridge ratio), EW/EFW (egg width/egg float width ratio).

| Parameter | T2 | IndCh | T6 | IndInt |
| --- | --- | --- | --- | --- |
| EL | 442.38±3.3 | 443.53±4.84 | 417.05±4.51 | 418.66±3.36 |
| EW | 134.7±2.12 | 150±2.92 | 128.82±1.89 | 130±2.18 |
| EL/EW | 2.96±0.06 | 3.30±0.06 | 3.21±0.04 | 3.22±0.05 |
| EFL | 218.09±3.49 | 229.41±4.81 | 210.58±6.21 | 209.33±10.62 |
| EFW | 64.76±1.48 | 71.17±6.96 | 67.64±7.52 | 69.33±7.98 |
| EFL/EFW | 3.40±0.09 | 3.24±0.08 | 3.14±0.12 | 3.15±0.15 |
| FR | 20.8±0.11 | 20.81±0.08 | 17.35±0.31 | 16.66±0.13 |
| EL/EFL | 2.04±0.03 | 2.0±0.07 | 2.13±0.11 | 1.97±0.06 |
| EFL/ER | 10.49±0.19 | 10.98±0.22 | 12.20±0.43 | 13.02±0.52 |
| EFL/EW | 1.46±0.03 | 1.7±0.03 | 1.64±0.05 | 1.61±0.06 |
| EFW/ER | 3.11±0.07 | 3.41±0.08 | 3.92±0.13 | 4.16±0.12 |
| EW/EFW | 2.34±0.07 | 1.90±0.04 | 1.92±0.05 | 1.93±0.08 |

There was no significant difference between the male and female mosquito wing length (Table 3) and between the parental line and their isofemale forms. The wing measurement values were found to be non-significant when compared between the isofemale lines of IndCh (F=1.32, p>0.05 for males; F=2.43, p>0.05 for females) and IndInt (F=1.31, p>0.05 for males; F=1.12, p>0.05 for females) and with their respective parental lines.

**Table 3:**
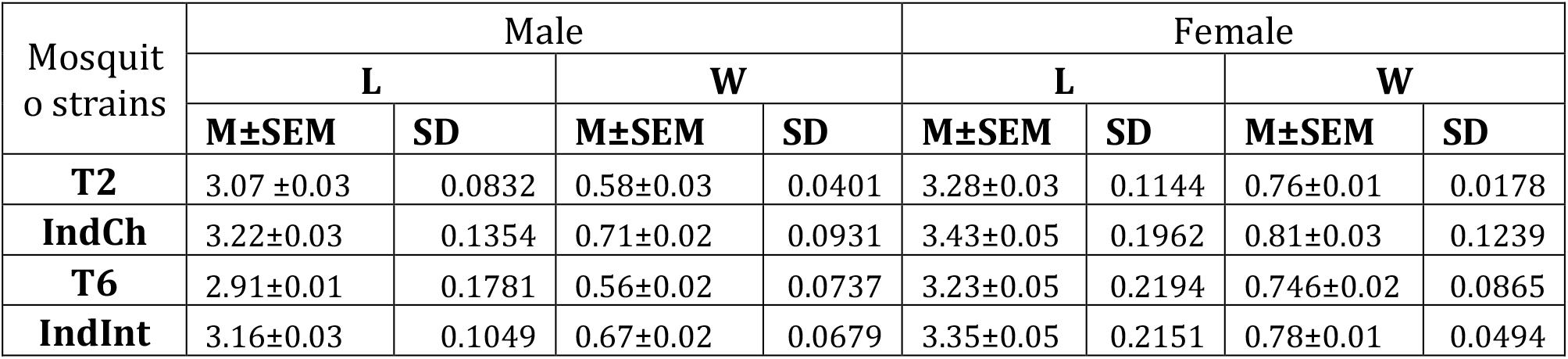
Wing measurement of paternal and isofemale strains of IndCh and IndInt variants.

| Mosquito strains | Male |  |  |  | Female |  |  |  |
| --- | --- | --- | --- | --- | --- | --- | --- | --- |
|  | L |  | W |  | L |  | W |  |
|  | M±SEM | SD | M±SEM | SD | M±SEM | SD | M±SEM | SD |
| **T2** | 3.07 ±0.03 | 0.0832 | 0.58±0.03 | 0.0401 | 3.28±0.03 | 0.1144 | 0.76±0.01 | 0.0178 |
| **IndCh** | 3.22±0.03 | 0.1354 | 0.71±0.02 | 0.0931 | 3.43±0.05 | 0.1962 | 0.81±0.03 | 0.1239 |
| **T6** | 2.91±0.01 | 0.1781 | 0.56±0.02 | 0.0737 | 3.23±0.05 | 0.2194 | 0.746±0.02 | 0.0865 |
| **IndInt** | 3.16±0.03 | 0.1049 | 0.67±0.02 | 0.0679 | 3.35±0.05 | 0.2151 | 0.78±0.01 | 0.0494 |

### Homozygosity achieved in isofemale lines

To decipher the levels of homozygosity achieved through sibling mating across filial generations, we obtained 100X sequence coverage using Illumina reads from DNA pooled from 50 individuals from the isofemale IndCh and IndInt strains homogenized for 5 and 14 generations, respectively. As a negative control, we used the dataset generated from a pool of 50+ individuals directly from a lab colony [40] of STE2 strain without genetic homogenization. The low ratios of heterozygous over homozygous SNPs seen for the two isofemale lines showed the decrease in heterozygosity achieved by isofemale lines around the 5th generation over pools of individuals from lab samples (Fig. 5).

**Figure 5:**
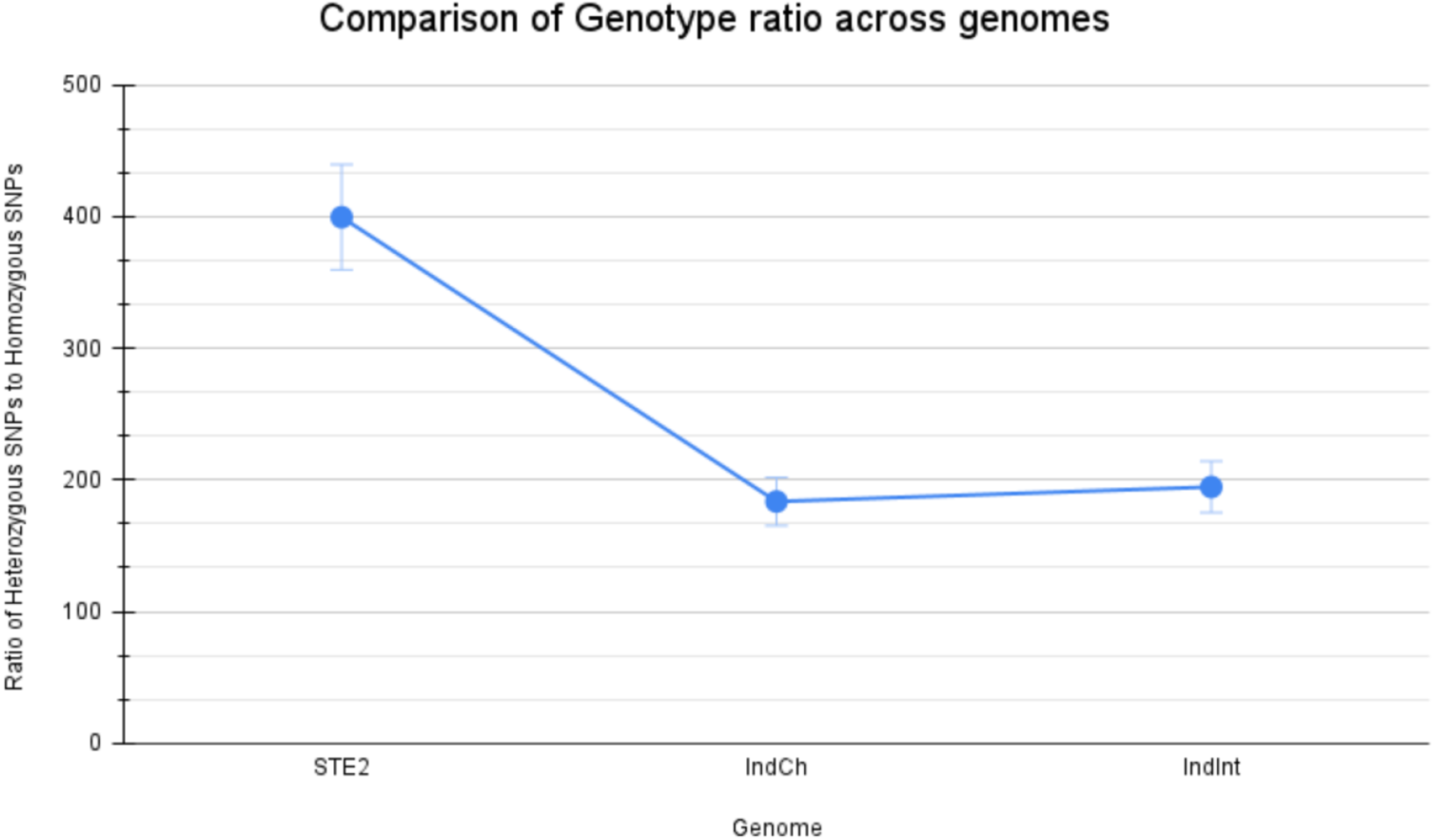
Ratio of heterozygous versus homozygous SNPs in the three strains including STE2 (not homogenized), IndCh (homogenized over 5 generations), and IndInt (homogenized over 14 generations).

It should be mentioned that while IndCh is homozygous for standard forms of all known inversions, STE2 is heterozygous for the 2R*b* genotype and IndInt is heterozygous for the 3L*i* genotype even after 14 generations of isofemale selection. Interestingly, the isofemale line for IndInt shows increased heterozygosity for 3L*i* relative to its parent line (unpublished data), perhaps amplifying the reduction in vectorial competence in the isofemale over its parental line (T6, Fig. 6 & 7). Even in the case of IndCh, the percentage of insects with the standard 2R*b* increased over generations of isofemale lines [78], perhaps explaining the increased vectorial competence of the isofemale line over that of the parental line (T2, Fig. 6 & 7).

**Figure 6:**
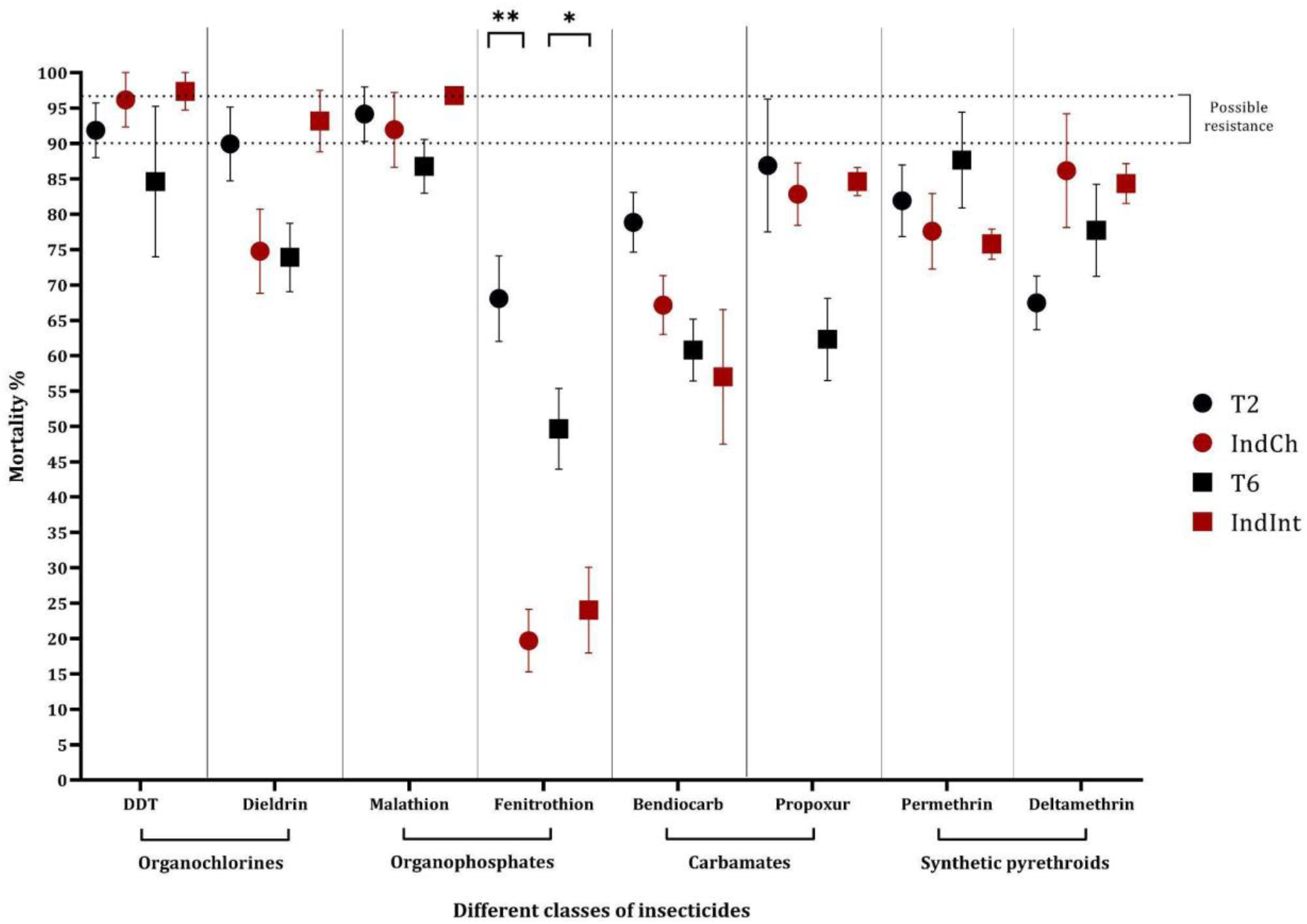
Summary plot of mortality percentage across 4 different classes of insecticides.

**Figure 7:**
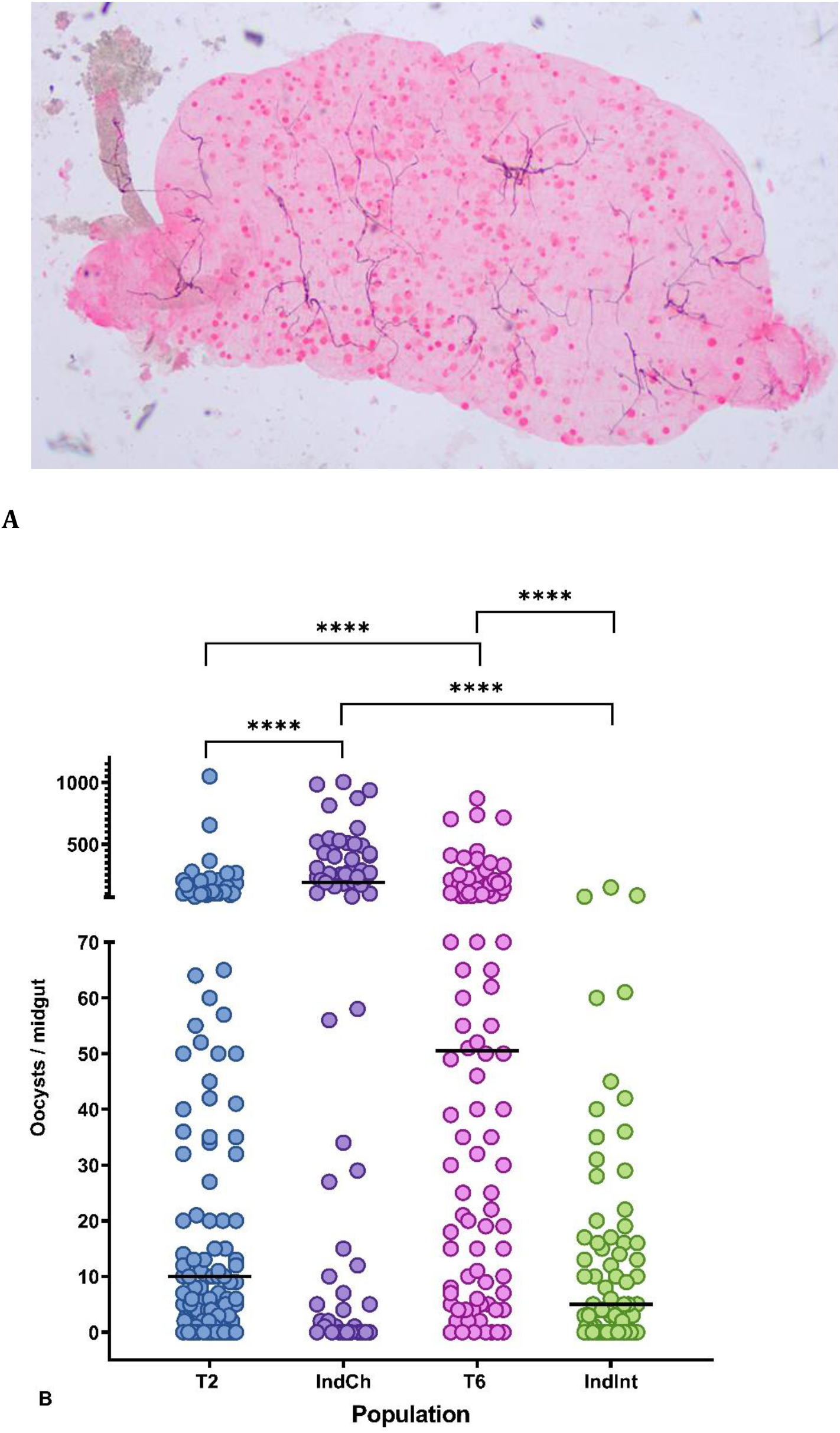
(A) Representative image of Pb-infected mosquito midgut. (B) Vectorial competence between the parental and isofemale lines for *Plasmodium berghei*. Each dot represents the number of oocysts in an individual midgut of mosquitoes, with median values (horizontal bar). Asterisks denote significant differences at p<0.0001 (Mann-Whitney non-parametric t-test).

### Susceptibility assay across four classes of insecticides

Insecticide susceptibility was determined for both isofemale and their parental colonies across four classes of insecticides [33]. The number of surviving individuals at 24 h post-exposure was calculated to understand the susceptibility/resistance status. The results suggest both the isofemale lines showed higher susceptibility to DDT than their parental lines. In case of dieldrin, IndCh showed a decreased mortality percentage of 71.6% than T2 which exhibits possible resistance. In contrast, the IndInt showed greater sensitivity (93% mortality) than T6 (68.2% mortality) as per WHO standards [60]. The change in the susceptibility might be due to different target sites of these two insecticides (Fig. 6). In Organophosphates, all the populations showed high mortality (>90%), except T6. For fenitrothion, both the isofemale lines exhibited significant resistance (<30% mortality), compared to their respective parental lines (non-parametric, Mann–Whitney test; p<0.0001). For Carbamate and Synthetic Pyrethroids, all the four mosquito colonies showed confirmed resistance (mortality <90%, ranging between 60% to 87%) (Fig. 6).

### Vectorial competence studies

#### P. berghei

Four colonies (T2, IndCh, T6, IndInt) were fed with *P. berghei* infected blood of BALB/c mice and evaluated for their infection rate. The infection rate was calculated for each colony as the percentage of mosquitoes with oocysts in their midguts. There was not much variation in mean infection rate of the type form T2 (82.73%) and IndCh (85.07%). However, it was interesting to note there was considerable difference in the mean infection rate between the intermediate form of T6 (92.86%) and IndInt (70.83%).

Among the four colonies, the highest oocyst range was observed in T2 (1-1050), followed by IndCh (1-1003), T6 (1-870) and IndInt (1-150). The mean number of oocysts per positive midgut in T2, IndCh, T6, and IndInt was 52.47 (SD±SEM=120.3±10.20), 235.6 (266.4±32.55), 112.4 (168.6±17.03), and 14.54 (24.39±2.87), respectively, whereas the median values were 10.00, 190.00, 50.50, 5.00 for T2, IndCh, T6, and IndInt, respectively. Furthermore, a comparison between parental colonies and isofemale colonies revealed a significant difference in their vector competence (p<0.0001, non-parametric, Mann–Whitney test, Supplementary-1).

#### P. falciparum

In the case of *P. falciparum*, there was not much variation in mean infection rate of the type forms, T2 (80.40%), and IndCh (78.57%) (Fig. 8B). The mean infection rate differed between the intermediate form of T6 (79.62%) and IndInt (69.57%). Among the four colonies, the highest oocyst range was observed in T6 (1-231), followed by IndInt (1-125), IndCh (1-106) and T2 (1-85). The mean number of oocysts per positive midgut in T2, IndCh, T6, and IndInt was 16.34 (SD±SEM=17.94±1.13), 26.39 (34.70±2.77), 18.39 (19.56±1.65), and 13.30 (19.63±1.67), respectively, whereas the median values were 12, 14, 16, 6 for T2, IndCh, T6, and IndInt, respectively.

**Figure 8:**
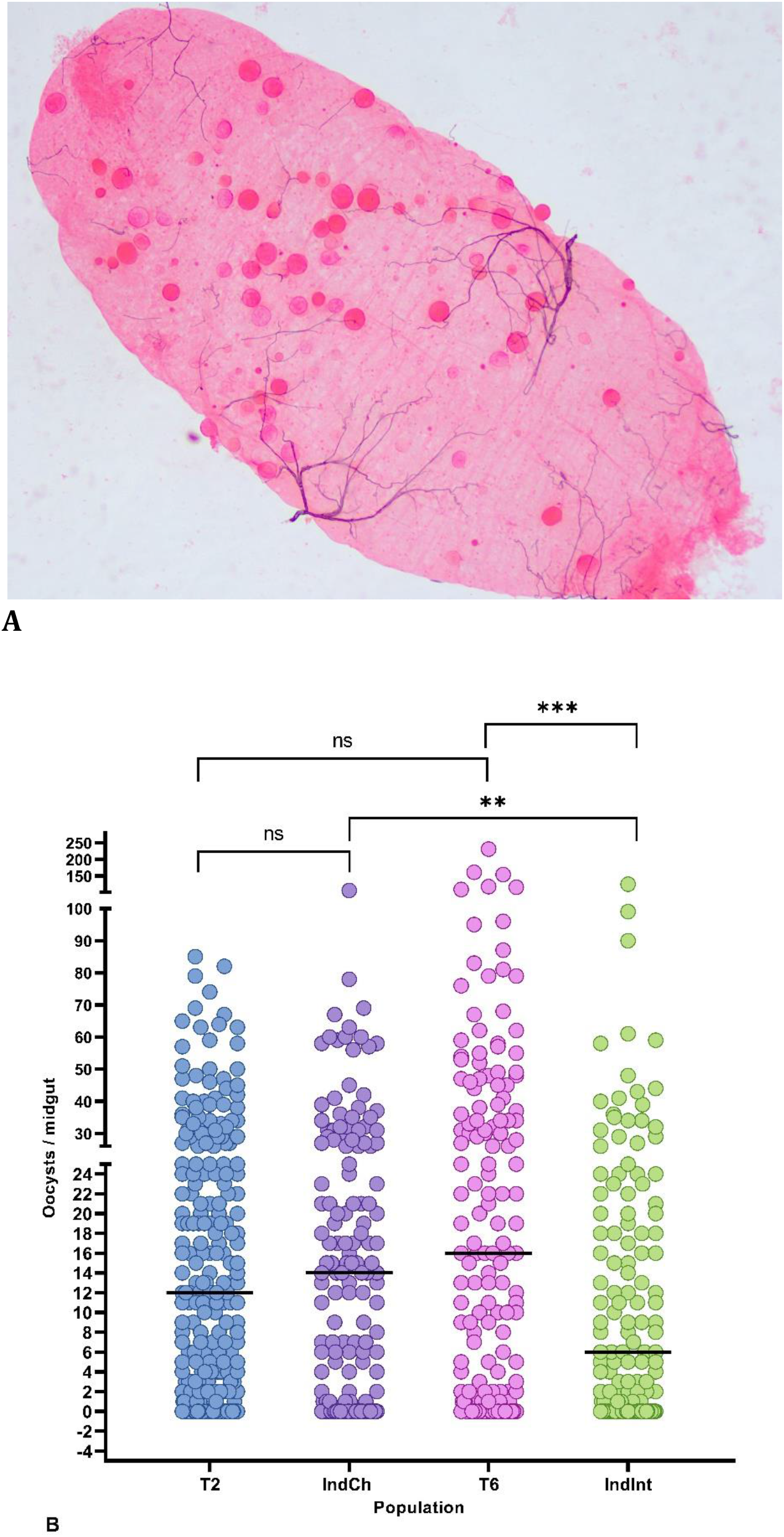
(A) Representative image of Pf-infected mosquito midgut. (B) Vectorial competence between the parental and isofemale lines for *Plasmodium falciparum*. Each dot represents the number of oocysts in an individual midgut of mosquitoes, with median values (horizontal bar). Asterisks denote significant differences at p<0.0001 (Mann-Whitney non-parametric t-test).

Further, a non-parametric Mann–Whitney comparative test between T2-T6 colonies (p<0.0563) and T2-IndCh colonies (p<0.3968) did not show any significant difference. However, the homozygous forms of the isofemale colonies, IndCh and IndInt showed significant difference (p<0.0045). Significant difference was observed between T6 vs IndInt (p<0.0003). These results corroborate earlier findings that the type form is a better vector when compared to the intermediate form [34, 41] (Fig. 8B).

## Discussion

The present study provides a detailed overview of the key considerations followed during the establishment of the isofemale line in the two biological forms of *An. stephensi*. We have meticulously characterized several life-cycle parameters in every generation during the creation of both isofemale lines to maintain their authenticity. During establishment, key emphasis was given to restore some characteristics of the parent population after the initial bottleneck to achieve normalization of fecundity, hatching, pupation, eclosion, and male: female ratio. Accordingly, we find that in both isofemale lines, homogeneity in these parameters is achieved within ~13 generations (Fig. 2).

While homogeneity in the biological parameters was achieved in ~13 generations, the impact of this on other traits, such as insecticide resistance and vectorial competence was dramatic. This is the first time a discrete study has been conducted by establishing two homozygous isofemale lines from the type and intermediate forms of *An. stephensi* and assessing their vectorial competence against two different malaria parasites, *P. berghei* and *P. falciparum*. The findings suggest that there is a significant enrichment in vectorial competence between the parental and corresponding isofemale lines. When mosquitoes were exposed to *P. berghei*-infected mice, there was increased susceptibility to the type form (IndCh) than to the intermediate form (IndInt). Furthermore, a significant difference in vectorial competence was seen between the parental and corresponding isofemale lines. This might be due to the high levels of homozygosity in certain alleles that might have enhanced the vectorial competence.

In the case of *P. falciparum-*infected blood meal, there was no significant difference between the parental and corresponding isofemale lines. However, there was a significant difference found between the two isofemale lines belonging to the type and intermediate forms, supporting earlier reports of the type form being a better vector than the intermediate form [25]. Establishing an isofemale line has been shown to delineate the non-additive genetic variance due to local adaptations, bottleneck events, and other epistatic effects [42]. We conclude that isofemale lines derived from type and intermediate forms achieve enrichment of characteristic phenotype with respect to vectorial competence expected for each form.

We also sequenced 100X coverage of DNA from a pool of 50 plus individuals from both IndCh and IndInt lines after 5 and 14 generations to assess the homogeneity achieved at the genome level. As a negative control, we used a public dataset with 100X coverage of STE2 strain where a pool of DNA from 50 individuals from a lab population. The ratio of the heterozygous versus homozygous variants in these reads relative to a gold-standard reference genome [43] clearly shows that genome level homozygosity is achieved within as few as 5 generations in IndCh. Further, the increased homozygosity in the organism provides an opportunity to decipher the genome architecture at a very high-resolution [43, 44]. While the high-resolution genome assembly of UCI and IndCh strains were achieved after 5 and 6 generations of isofemale lines, the IndInt line was homogenized over 14 generations before attempting an assembly because, as shown in Figure 2a, increased generations were needed to normalize fecundity. This could be because unlike UCI and IndCh with no heterozygous paracentric inversions, IndInt retains a large paracentric inversion even after 14 generations. Furthermore, our data show (unpublished data) that the percentage of this inversion increased considerably from that of the parent population. This may be a notable example of the complexity of maintaining a given trait while retaining fitness.

In the present study, we observed significant variations in the founding population and its isofemale progeny. This might be a result of the segregation of the genotypes resulting in the better demarcation of the susceptible/resistant phenotype in comparison to the parental lines. This can be further studied for a better understanding of underlying mechanisms of resistance, which is relatively easier to identify in isofemale lines due to their homozygosity.

The two isofemale lines showed various levels of susceptibility to each class of insecticide. In all the four colonies including the two parent and two isofemale lines, the levels of resistance varied across the two insecticides belonging to the same class, such as DDT and Dieldrin belonging to the class Organochlorines. For Dieldrin, the parental line T2 exhibited greater sensitivity than its isofemale IndCh, whereas the parental line T6 showed greater resistance than its isofemale form IndInt. The difference in the mode of action of these two insecticides might be the reason for this trend. For example, DDT resistance is associated with mutation(s) in the target site of the voltage-gated sodium channel (vgsc) gene and metabolic resistance [45, 46], whereas dieldrin resistance is due to a single point mutation in GABA receptors [47]. Also, among the Organophosphates, resistance was observed to fenitrothion, but susceptibility was seen for malathion among all the four colonies. For fenitrothion, there was significant difference in the susceptibility status between the parental and their respective isofemale lines. Earlier studies had suggested that many strains of *An. stephensi* collected from different parts of India were susceptible to fenitrothion [29]. However, for carbamates and synthetic pyrethroids, all the four populations showed resistance to both class of the insecticides. In view of the fact that the insectary-colonized mosquitoes and their isofemale lines were not under any selection pressure from insecticides and were grown without any insecticide exposure, they continued to maintain this resistance.

The observed variations in insecticide susceptibility might be due to large structural variants such as inversions. Recent advances have revealed the possible role of structural variations like duplications and inversions in insecticide resistance, which would be interesting to dissect among isofemale colonies in the future [48]. The role of structural variation has also been shown implicated in different mechanisms of copy number variation, transposable element insertions, and tandem duplication of different metabolic genes [49, 50]. Characterization of the isofemale lines during their establishment as reference genomes revealed that there is a homozygous standard 2R form in the IndCh isofemale line [44] and a heterozygous 3L*i* inversion in the IndInt isofemale line (unpublished data). The functional characterization of the 3L*i* region revealed 36 isoforms of cuticle-forming genes in the IndInt isofemale line. Earlier studies suggest that cuticle thickening is associated with pyrethroid and insecticide resistance in other *Anopheles* species [51]. Similarly, the 2R*b* region has 1353 predicted genes, which include several genes associated with insecticide resistance such as ACE1, as well as tandem clusters of GST and Cyp450 paralogs in the IndCh isofemale line [44].

### Variations in vector competence among the biological forms

Although the various biological forms can be clearly delineated using the number of ridges on the egg float, there have been efforts to develop molecular markers to identify the various forms. These include mitochondrial genes cytochrome oxidase 1 (COI) and cytochrome oxidase 2 (COII) [52], the rDNA-ITS2 and domain-3 (D3) marker [53], Odorant-binding protein *Anste*Obp1 marker [54] and microsatellite markers [55] to characterize the different biological forms. A single SNP has been previously reported [54] among the various forms of *An. stephensi* found in Iran. When we tested the validity of this SNP for Indian *An. stephensi* populations (insectary-colonized and wild populations) that were collected from 5 different locations, we found that most of the individuals from the wild had the SNP (G) in *AsteObp1* (intron I Position 91bp) reported to be diagnostic of the type form, 18/105 individuals had an A at this same position, 18/105 had an A/G and 1/105 had an A/C in that position, likely representing population-level variations that have not been reported earlier (unpublished data). Also, many of the other markers are not universal and may vary from region to region and cannot resolve the biological forms accurately [56]. Here, we clearly demonstrate that the number of egg ridges is the gold standard and applicable marker to develop isofemale lines.

There are also reports of a third form with the lowest egg-float ridge number that is known as ‘var. *mysorensis’* [25, 31, 34, 57], but this has not been established or characterized in the current study. The *mysorensis* form is mostly found in rural areas and breeds in fresh water sources like wells, ponds, cement tanks, streams, etc. Earlier studies have shown var. *mysorensis* to exhibit variations in their insecticide susceptibility status [58] and vectorial competence [41]. The *mysorensis* form is highly zoophilic and has very poor vectorial competence. A thorough characterization of this third biological form might reveal cues that can be exploited in vector management strategies.

Overall, the findings of this paper provide a benefit of developing isofemale lines from the insectary-colonized mosquito populations. Further, we showed that as low as 5 filial generations of sibling mating may be sufficient in *An. stephensi* to achieve greater levels of homozygosity. Greater number of sibling mating may be required (>13 generations) for establishing isofemale lines with optimal fitness. The isofemale lines thus developed could be used to elucidate mechanisms underlying insecticide resistance, which is more pronounced than in the parental lines. The isofemale lines should also prove useful in understanding the basis for the greater vectorial competence of the type variant, relative to the intermediate form for both *P. berghei* and *P. falciparum*. By assembling high-quality reference genomes for IndInt and IndCh we have been able to hypothesize genetic factor/structural variant behind the differential vectorial competence [59].

## Author contributions

CG: Established isofemale lines, recorded life-table data, egg morphometric study, and *P. berghei* vectorial competence assay.

NK: *P. falciparum* vectorial competence assays, analyzed the data and insectary-related work.

RBSK: Insecticide susceptibility assays.

SM, SGJ, CK: Insectary-related work, and isofemale line maintenance.

TA: Bioinformatic analysis.

SSri: Overseeing bioinformatics work, helping in reviewing the manuscript draft.

SS: Conceptualization, funding, and contributions to the manuscript.

SK: Conceptualizing and writing the first draft, analyzing the data, and insectary-related work.

SunS: Coordinating and overseeing the entire insect work and helping in reviewing the manuscript draft.

All authors read and approved the final manuscript.

## Acknowledgment

The authors thank Dr. Rakesh Mishra, Director of the Tata Institute for Genetics and Society (TIGS), Bengaluru, India, for his support. We are thankful to Dr. Sushanta Kumar Ghosh, Senior Scientist (retd.) of ICMR-NIMR, Bengaluru, for guiding the establishment of TIGS insectary. We are grateful to Dr. Kailash Patra, R&D Director at Intrivo Diagnostics, California, United States for giving training on parasite culture. We are thankful to Mr. Joydeep Roy, TIGS for his technical support. Animal work in the NCBS/inStem Animal Care and Resource Center was partially supported by the National Mouse Research Resource (NaMoR) grant #BT/PR5981/MED/31/181/2012;2013-2016 & 102/IFD/SAN/5003/2017-2018 from the Department of Biotechnology, GoI. The authors thank Tata Institute for Genetics and Society (TIGS) India for funding researchers involved in this work and IBAB for bioinformatics work.

